# Spontaneous thalamic activity modulates the cortical innervation of the primary visual nucleus of the thalamus

**DOI:** 10.1101/2020.11.13.382366

**Authors:** Verónica Moreno-Juan, Mar Aníbal-Martínez, Álvaro Herrero-Navarro, Miguel Valdeolmillos, Francisco J. Martini, Guillermina López-Bendito

## Abstract

Sensory processing relies on the correct development of thalamocortical loops. Visual corticothalamic axons (CTAs) invade the dorsolateral geniculate nucleus (dLGN) of the thalamus in early postnatal mice according to a regulated program that includes activity-dependent mechanisms. Spontaneous retinal activity influences the thalamic incursion of CTAs, yet the perinatal thalamus also generates intrinsic patterns of spontaneous activity whose role in modulating afferent connectivity remains unknown. Here, we found that patterned spontaneous activity in the dLGN contributes to proper spatial and temporal innervation of CTAs. Disrupting patterned spontaneous activity in the dLGN delays corticogeniculate innervation under normal conditions and upon eye enucleation. The delayed innervation was evident throughout the first two postnatal weeks but resumes after eye-opening, suggesting that visual experience is necessary for the homeostatic recovery of corticogeniculate innervation.

## Introduction

Visual information flows from the retina to the primary visual cortex (V1) relayed by the primary visual nucleus of the thalamus, the dorsal lateral geniculate nucleus (dLGN). However, the visual message arriving at V1 contains information provided not only by the retina but also by other brain structures. In the dLGN, visual information from the retina is modulated by signals coming from the superior colliculus, thalamic reticular nucleus, brainstem, and feedback projections from layer 6 (L6) neurons of V1 (Guido, 2018). This corticothalamic loop is one of the largest dLGN inputs (Erişir et al., 1997) and seems to influence spike timing, structure of receptive fields or the amplitude of the response in immature and mature stages (Jurgens et al., 2012; Petrof and Sherman, 2013; Denman and Contreras, 2015; Murata and Colonnese, 2016; Hasse and Briggs, 2017). Thus, the arrival of relevant information at the visual cortex is likely to depend on the correct assembly and function of the corticothalamic connection between V1 and the dLGN.

The assembly of synaptic contacts between dLGN cells and both retinal and cortical projections follow a tightly regulated spatial and temporal organization. In mice, retinal axons start to innervate the dLGN before birth and, by postnatal day 2 (P2), they cover the entire nucleus (Godement et al., 1984). On the other hand, cortical axons target the ventral edge of the dLGN at P0 but they do not innervate the nucleus during the first postnatal days. Cortical axons start to innervate the dLGN at P3, occupying the entire nucleus by the end of the second postnatal week (Jacobs et al., 2007). Thus, although corticothalamic axons (CTAs) arrive when retinothalamic axons (RTAs) begin to innervate the dLGN, CTAs do not enter the dLGN until RTAs have spread throughout the nucleus.

The temporal dynamics of CTA innervation of the dLGN suggests that RTAs could orchestrate this process. Indeed, the absence or premature entry of RTAs into the dLGN accelerates corticogeniculate innervation (Seabrook et al., 2013). However, we still ignore the nature of the mechanism involved in this interaction between retinal and cortical inputs. One possibility is that RTAs secrete or control the secretion of molecular signals that modify the progression of CTAs. For example, the retinal input regulates the synthesis of aggrecan, a repulsive molecule that is present in neonatal dLGN and downregulated at the moment of corticogeniculate innervation (Brooks et al., 2013).

Apart from the trophic effects exerted by molecular signaling, activity-dependent mechanisms could be implicated in regulating innervation in the visual system. For example, CTAs accelerate their entry into the dLGN when retinal activity is disrupted in postnatal mice (Grant et al., 2016). Also, patterns of activity generated in the thalamus might influence corticogeniculate projections. Indeed, we have recently shown that the perinatal thalamus generates an intrinsic pattern of spontaneous activity that is known to regulate the organization of sensory systems (Moreno-Juan et al., 2017; Antón-Bolaños et al., 2019). It is possible that intrinsic patterns of thalamic activity regulate the ingrowth of CTA to dLGN. Here, we found that disrupting the pattern of spontaneous activity delayed the entry of CTAs into the dLGN, a delay that is promptly recovered by visual experience.

## Results

### Disrupting the pattern of spontaneous activity in the developing visual thalamus

To determine whether innervation of the dLGN by visual CTAs requires spontaneous thalamic activity, we disrupted the activity of the thalamus by overexpressing the inward rectifier potassium channel 2.1 (*Kcnj2*) in all principal thalamic sensory nuclei. As such, we administered tamoxifen to pregnant *Gbx2^CreER^; Rosa26 ^fs-Kcnj2^* mice (referred herein as *Th^Kir^*) on gestational day 10.5. Although *Gbx2* is expressed in amacrine cells of the retina at later stages (Kerstein et al., 2020), at this early stage of embryogenesis, the overexpression of Kir2.1 (fused to mCherry) was induced specifically in thalamic neurons (**Figure S1**) (Antón-Bolaños et al., 2019). We obtained the activity pattern of the dLGN using brain slices from control and *Th^Kir^* mice at P0 loaded with a fluorescent calcium indicator. We subdivided the dLGN area into regions of interest (ROIs) and measured the changes in fluorescence from each ROI (**Figure 1A**). The different ROIs became activated asynchronously or as part of synchronous ensembles of different sizes (**Figure 1B-1C**). Synchronous calcium events were grouped into two different categories: large propagating ensembles (waves) engaging almost the whole dLGN or relatively small ensembles engaging less than 50% of the nucleus. In control slices, synchronous events encompassed small and large ensembles. In contrast, synchronous calcium events in the *Th^Kir^* only included small ensembles (**Figure 1D**) and the overall activity was diminished (**Figure 1E**). Therefore, and as shown previously for the somatosensory and auditory nuclei of the thalamus (Moreno-Juan et al., 2017; Antón-Bolaños et al., 2019), overexpression of Kir2.1 in the dLGN reduced the overall frequency of spontaneous calcium transients and switched the structure of synchronous events from large ensembles to mostly small ensembles.

**Figure 1.**
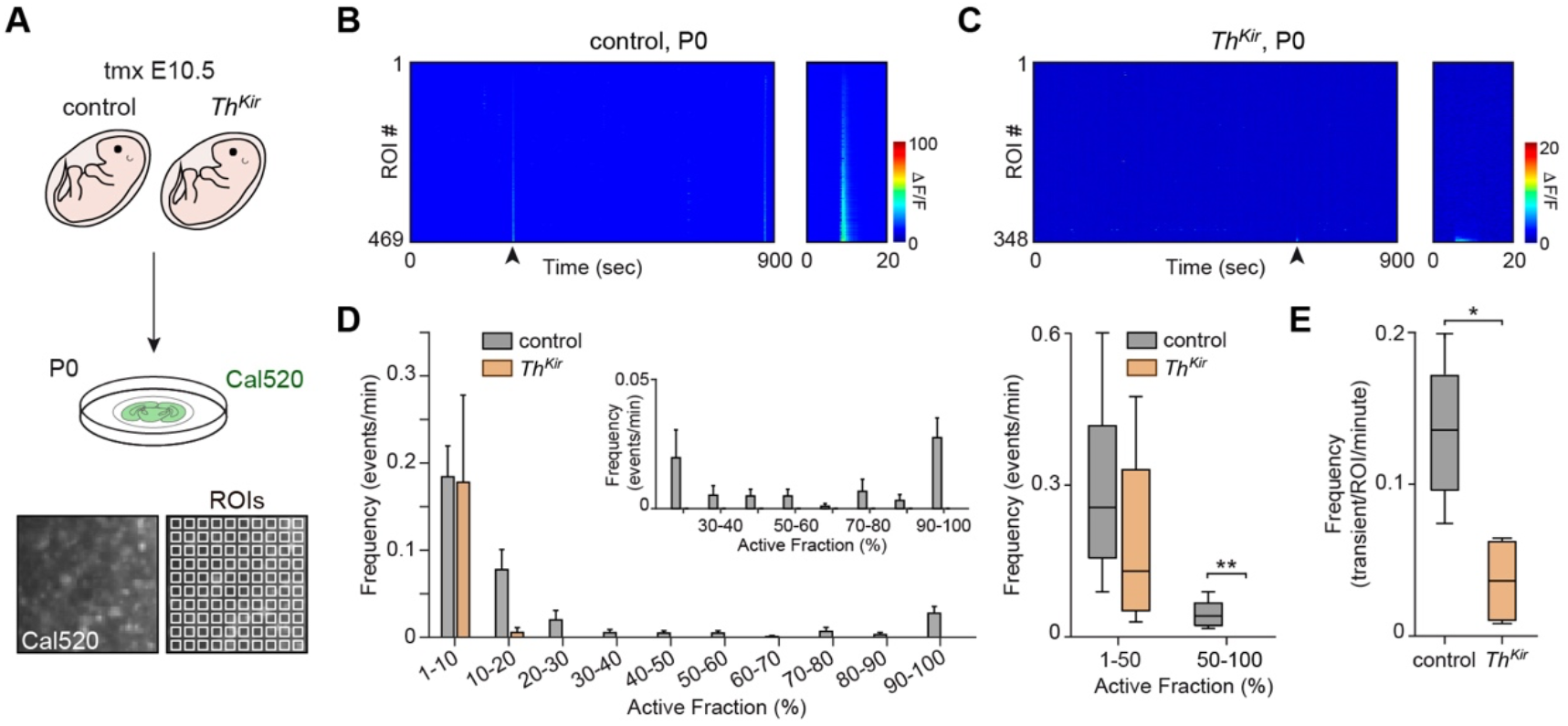
Overexpression of Kir2.1 in the dLGN disrupts the pattern of spontaneous activity. (A) Scheme of the experimental design used for calcium imaging in brain slices from control and *Th^Kir^* mice at P0. At the bottom, pictures of a small zone of the dLGN and the overlaying grid of regions of interest (ROIs). (B) Example of the calcium signal obtained from all dLGN ROIs in a 15-minute recording from a control slice. The arrowhead points to an event of synchronous activation of multiple ROIs. This event comprises almost 100% of the ROIs and is shown at a larger temporal magnification on the right. Another large event has been detected at the end of the recording. (C) Example of the calcium signal obtained from all dLGN ROIs in a 15-minute recording from a *Th^Kir^* slice. The arrowhead points to an event of synchronous activation of multiple ROIs. This event comprises less than 10% of the ROIs and is shown at a larger temporal magnification on the right. (D) Distribution of mean frequencies of synchronic calcium events that activate more than 1% of the ROIs in the dLGN of control and *Th^Kir^* mice. The inset shows the same information for synchronic events that activate more than 20% of the ROIs. On the right, mean frequency of the synchronic events grouped by active fraction into two categories (1-50% and 50-100%) in the dLGN of control and *Th^Kir^* slices at P0 (control *n* = 10, *Th^Kir^ n* = 4). (E) Mean frequency of calcium transients in dLGN ROI of control and *Th^Kir^* slices (control *n* = 10, *Th^Kir^ n* = 4). Graphs represent mean ± SEM, each dot corresponding to a single experimental unit (1 slice per mouse): *P<0.05, **P<0.01, ***P<0.001. Scale bars, 100 μm.

### Thalamic spontaneous activity is involved in the entry of CTAs into the dLGN

In light of the above, we set out to determine whether an abnormal intrinsic pattern of spontaneous activity in the thalamus might affect the progression of CTAs into the dLGN. As such, we visualized CTAs in the dLGN by immunostaining against vGlut1 transporter which is expressed at the terminals of corticothalamic neurons or by using the Golli-tau-eGfp mouse that labels L6 and a subpopulation of L5 cortical neurons (Jacobs et al., 2007; Piñon et al., 2009). L5 projections target the lateral posterior nucleus and do not normally arborize within the dLGN. Spontaneous activity did not affect the initial targeting of CTAs since L6 axons were located at the border of the dLGN in both control and *Th^Kir^* mice at P0 (**Figure 2A**). At P4, CTAs had already entered the dLGN of control mice, while they remained at the border in *Th^Kir^* mice (**Figure 2B**, left panels, and **Figure 2C**). Between P6 and P9, CTAs covered more than half the nucleus in control mice, however, *Th^Kir^* CTAs covered less than 20% of the nucleus (**Figure 2B**, right panels, and **Figure 2C**). We further confirm the visual origin of the delayed CTAs in the *Th^Kir^* model by infecting V1 with a lentivirus expressing *Gfp* at P3. We processed the brains at P9 and found fewer CTAs in the dLGN of *Th^Kir^* than in control mice, as in VGlut1 immunostainings (**Figure 2D**). Together these results indicate that the innervation of the dLGN by visual CTAs is delayed when thalamic spontaneous activity is disrupted.

**Figure 2.**
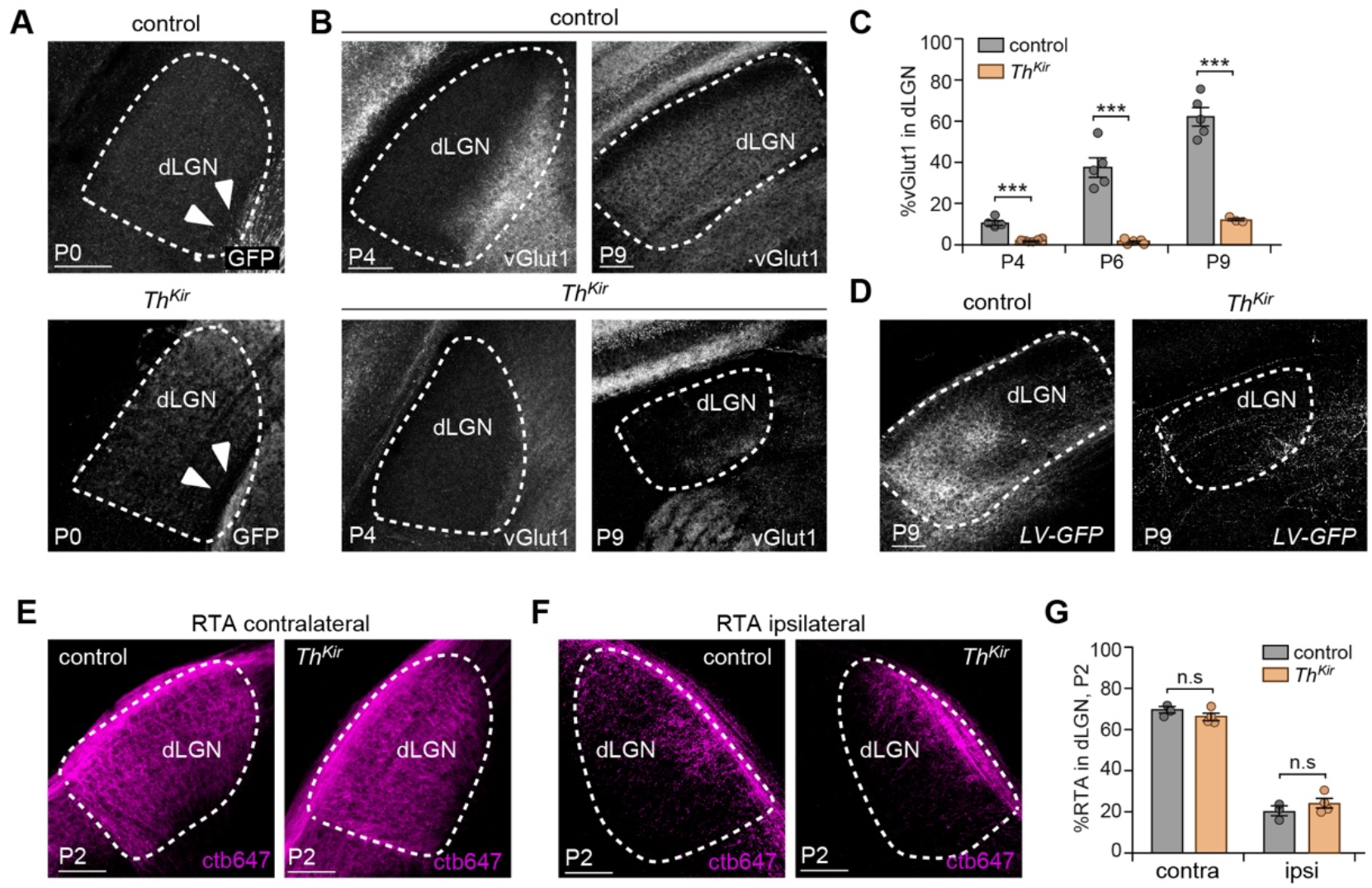
Corticogeniculate innervation is delayed in *Th^Kir^* mice. (A) Coronal sections of Golli-tau-eGfp control and *Th^Kir^* brains at P0 showing the expression of GFP in CTAs (arrowheads) in the dLGN (control *n* = 3, *Th^Kir^ n* = 6). (B) Coronal sections of control and *Th^Kir^* brains at P4 and P9 showing vGlut1 expression in CTAs. (C) Quantification of the relative dLGN area occupied by vGlut1 in control and *Th^Kir^* slices at different postnatal stages (P4 control *n* = 5, *Th^Kir^ n* = 5; P6 control *n* = 5, *Th^Kir^ n* = 5; P9 control *n* = 5, *Th^Kir^ n* = 3). (D) Coronal sections of control and *Th^Kir^* P9 brains that were injected with a lentivirus coding for *Gfp* into the V1. (E) CTB-647 labelling of the contralateral retinal axons in the dLGN of control and *Th^Kir^* coronal slices at P2. (F) CTB-647 labelling of ipsilateral retinal axons in the dLGN of control and *Th^Kir^* coronal slices at P2. (G) Quantification of the relative dLGN area occupied by contralateral and ipsilateral RTAs at P2 (control *n* = 3, *Th^Kir^ n* = 4). Graphs represent mean ± SEM, each dot corresponding to a single experimental unit (mouse): *P<0.05, **P<0.01, ***P<0.001, n.s. not significant. Scale bars, 100 μm.

In addition to this role of thalamic activity demonstrated above, corticogeniculate innervation of the dLGN is controlled by retinal input (Brooks et al., 2013; Seabrook et al., 2013; Grant et al., 2016). Retinal projections arborized throughout the dLGN, occupying 90% of its area at P2, when CTAs start to invade the nucleus (Godement et al., 1984; Jacobs et al., 2007; Seabrook et al., 2013). Thus, we tested whether an abnormal ingrowth of RTAs into the dLGN in *Th^Kir^* mice might underlie delayed CTAs. To this end, we labeled retinal projections by injecting cholera toxin subunit B into the retina. We found that contralateral and ipsilateral RTA innervation of the dLGN was normal in *Th^Kir^* mice at P2 (**Figure 2E-2G**). Hence, RTAs targeted and invaded the dLGN independently of thalamic activity and therefore, the delayed invasion of CTAs into the dLGN in *Th^Kir^* mice is not due to a defect in RTA innervation.

### Disrupting spontaneous activity in the thalamus leads to dLGN neuron death

We observed alterations in dLGN shape when thalamic spontaneous activity was disrupted. While from P0 to P3, the size of the dLGN was comparable to the control, it remained static from P4 onwards, coinciding with the initial phase of CTA innervation (**Figure S2A**). To determine whether this lack of growth of the dLGN might be due to an increase in cell-death or the enhanced compaction of dLGN cells, we analyzed caspase-3a activity (Ahern et al., 2013; Lossi et al., 2014) and cell density in the dLGN of control and *Th^Kir^* mice. There was a significant increase in caspase-3a immunostaining from P3 to P6 in the dLGN of the *Th^Kir^* mice (**Figure S2B** and **S2C**) but no increase in cell density (**Figure S2D** and **S2E**). Hence, the diminished size of the dLGN in *Th^Kir^* mice might be due to a higher rate of cell-death, which may in turn be related to the absence of CTAs or to abnormal spontaneous activity (Blanquie et al., 2017).

### Retinal inputs modify the spontaneous thalamic activity and regulate the innervation of the dLGN by CTAs

The impact of RTAs on the entry of CTAs into the dLGN is evident upon perturbing the normal development of retinal projections. CTAs enter into the dLGN prematurely when retinal projections are genetically ablated or removed by neonatal enucleation, as well as, when the pattern of retinal activity is abnormal (Brooks et al., 2013; Seabrook et al., 2013; Grant et al., 2016). We corroborated these data using an experimental model (called embBE) where both eyes were enucleated at embryonic day (E)14, before retinal axons arrive at the dLGN (**Figure 3A**). Whereas CTAs in control mice recapitulated the previously described developmental invasion of the dLGN, CTAs in embBE mice covered a significantly higher proportion of the nucleus at P2. This acceleration of the dLGN invasion was maintained at least until P7, the last developmental stage studied here (**Figure 3B**, **3C,** and **Figure S3**).

**Figure 3.**
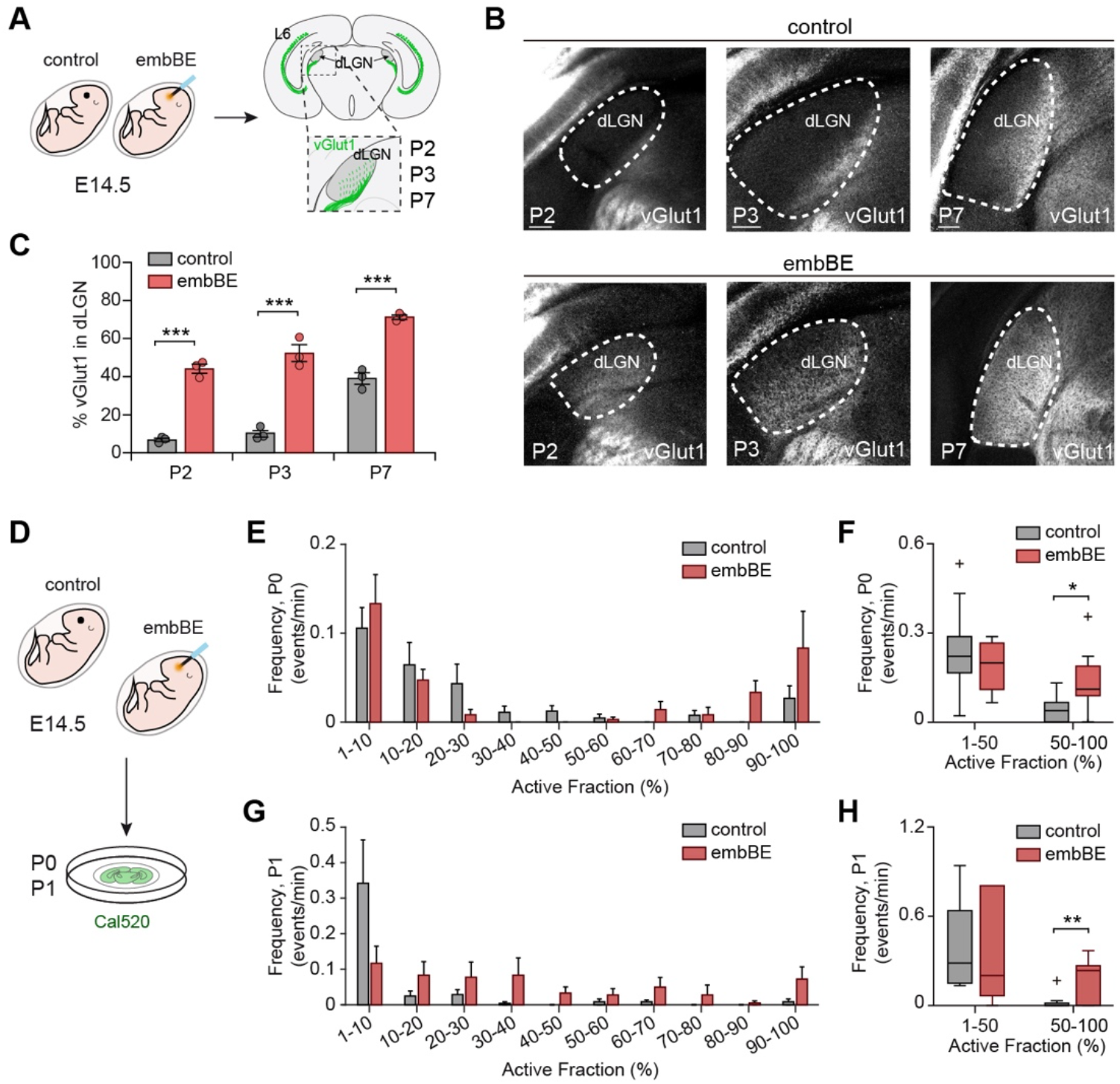
Retinal input modulates cortical innervation and the pattern of spontaneous activity in the dLGN. (A) Scheme of the experimental design used to stain cortical axons in pups that were enucleated at E14.5 and control littermates. (B) Coronal brain sections of control and embBE mice showing vGlut1 immunostaining of CTAs in the dLGN at P2, P3 and P7. (C) Quantification of the relative dLGN area occupied by vGlut1-positive axons (P2: control *n* = 3, embBE *n* = 3; P3: control *n* = 3, embBE *n* = 3; and P7: control *n* = 3, embBE *n* = 3). (D) Scheme of the experimental design used for *ex vivo* calcium imaging of the dLGN in control and embBE mice at P0 and P1. (E) Distribution of mean frequencies of synchronic calcium events that activate more than 1% of the ROIs in the dLGN of control and embBE mice at P0. (F) Mean frequency of the synchronic events grouped by active fraction into two categories (1-50% and 50-100%) in the dLGN of control and embBE mice at P0 (control *n* = 9, embBE *n* = 7). (G) Distribution of mean frequencies of synchronic calcium events that activate more than 1% of the ROIs in the dLGN of control and embBE mice at P1. (H) Mean frequency of the synchronic events grouped by active fraction into two categories (1-50% and 50-100%) in the dLGN of control and embBE mice at P1 (control *n* = 8, embBE *n* = 6). Graphs represent mean ± SEM, each dot corresponding to a single experimental unit (mouse): *P<0.05, **P<0.01, ***P<0.001. Scale bars, 100 μm. The plus signs indicate outlier values.

It is likely that retinal projections influence the dynamics of corticogeniculate projections via trophic factors and the effect of retinal activity (Brooks et al., 2013). However, here, we also considered another tentative mechanism. In light of our previous findings, we cannot rule out that the absence of retinal input perturbs spontaneous activity in the dLGN, and that these abnormal patterns may contribute to the defects observed in corticothalamic innervation. To start testing this possibility, we recorded postnatal spontaneous activity in embBE mice using calcium imaging in slices. At P0, dLGN from embBE mice exhibited a higher frequency of large propagating waves compared with control mice (**Figure 3D-F**). Moreover, while large propagating waves normally vanished by P1 in control littermates, they remained in embBE mice, overlapping with the initial phase of the premature entry of CTAs into the dLGN (**Figure 3G** and **3H**). Thus, the pattern of spontaneous activity in the dLGN is not normal when retinal input is not present. However, in contrast to the *Th^Kir^* model, where the absence of large propagating events was associated with a delayed entry of CTAs, an increased frequency of large propagating events in the embBE model was associated with a premature corticogeniculate innervation.

Next, we wondered what the contribution was of thalamic spontaneous activity to the accelerated entry of CTAs into the dLGN in the absence of retinal input. To address this question, we suppressed large events of synchronous activity from the dLGN of embBE mice by performing bilateral embryonic enucleation in *Th^Kir^* mice (**Figure 4A-4C**). In this scenario, we found that the CTAs were accelerated but to a lesser extent than in embBE mice (**Figure 4D-4F**). Together these results corroborated the strong influence of RTAs over CTAs and showed that the temporal dynamics of CTA entry into the dLGN partly relies on its pattern of spontaneous activity.

**Figure 4.**
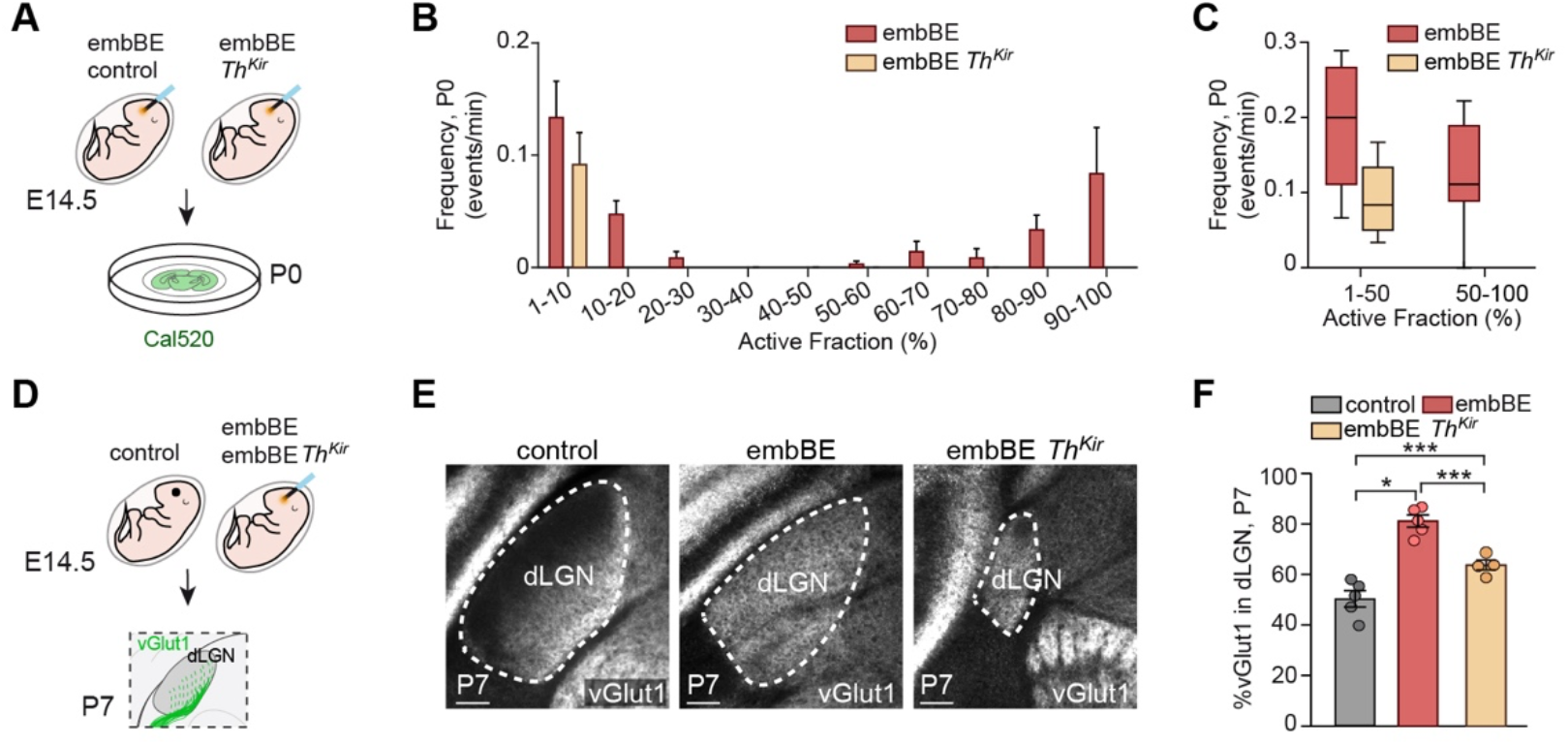
Disrupting spontaneous thalamic activity in embBE mice delays the entry of CTAs into the dLGN. (A) Scheme of the experimental design used to acquire spontaneous activity in the dLGN of embBE and embBE *Th^Kir^* mice. Tamoxifen was administered at E10.5 and enucleation was performed at E14.5 in control and *Th^Kir^* mice. (B) Distribution of mean frequencies of synchronic calcium events that activate more than 1% of the ROIs in the dLGN of embBE and embBE *Th^Kir^* mice at P0. (C) Mean frequency of the synchronic events grouped by active fraction into two categories (1-50% and 50-100%) in the dLGN of embBE and embBE ThKir mice at P0 (embBE *n* = 7, embBE *Th^Kir^ n* = 4). (D) Scheme of the experimental design used to reveal CTAs in control embBE and embBE *Th^Kir^* mice. Tamoxifen was administered at E10.5 and enucleation was performed at E14.5 in control and *Th^Kir^* mice. (E) Coronal sections showing vGlut1 immunostaining in the dLGN of embBE (*n* = 5), embBE *Th^Kir^* (*n* = 4), and control (*n* = 5) mice at P7. (F) Quantification of the relative dLGN area occupied by vGlut1-positive axons in embBE, embBE *Th^Kir^*, and control mice at P7. Graphs represent mean ± SEM, each dot corresponding to a single experimental unit (mouse): *P<0.05, **P<0.01, ***P<0.001. Scale bars, 100 μm.

### Visual experience rescues CTA innervation of the dLGN in *Th^Kir^* mice

Our results demonstrated that altering early patterns of spontaneous thalamic activity affects how cortical connections to the dLGN develop. Disrupting spontaneous activity in the dLGN delayed CTA innervation at least until P9 (**Figure 2B-D**). Since circuit remodeling in the visual system continues beyond the second postnatal week in mice (Hensch, 2005; Hooks and Chen, 2020), we questioned whether the visual pathway defects of the *Th^Kir^* model might persist in adult mice. Notably, at P30 CTAs had innervated the full extent of the dLGN in *Th^Kir^* mice as in control littermates (**Figure S4**), indicating that aberrant CTA innervation detected early in development is overcome at later stages.

Eye-opening occurs in mice at around P13, an event that is crucial for the final maturation and refinement of the visual system (Rochefort et al., 2009; Ko et al., 2013; Riyahi et al., 2021). As the decreased corticothalamic innervation of the dLGN observed at P9 is rescued by P30, we assessed whether eye-opening might play a role in this recovery. We analyzed the innervation of the dLGN by CTAs in both control and *Th^Kir^* mice at P10, P13 and P15, just before and after eye-opening. At P10, a massive invasion of CTAs into control dLGN was evident, while few CTAs were found in *Th^Kir^* dLGN (**Figure 5A** and **5C**). However, after eye-opening, we observed that the profiles of CTA invasion in control and *Th^Kir^* mice were indistinguishable (**Figure 5B** and **5D**).

**Figure 5.**
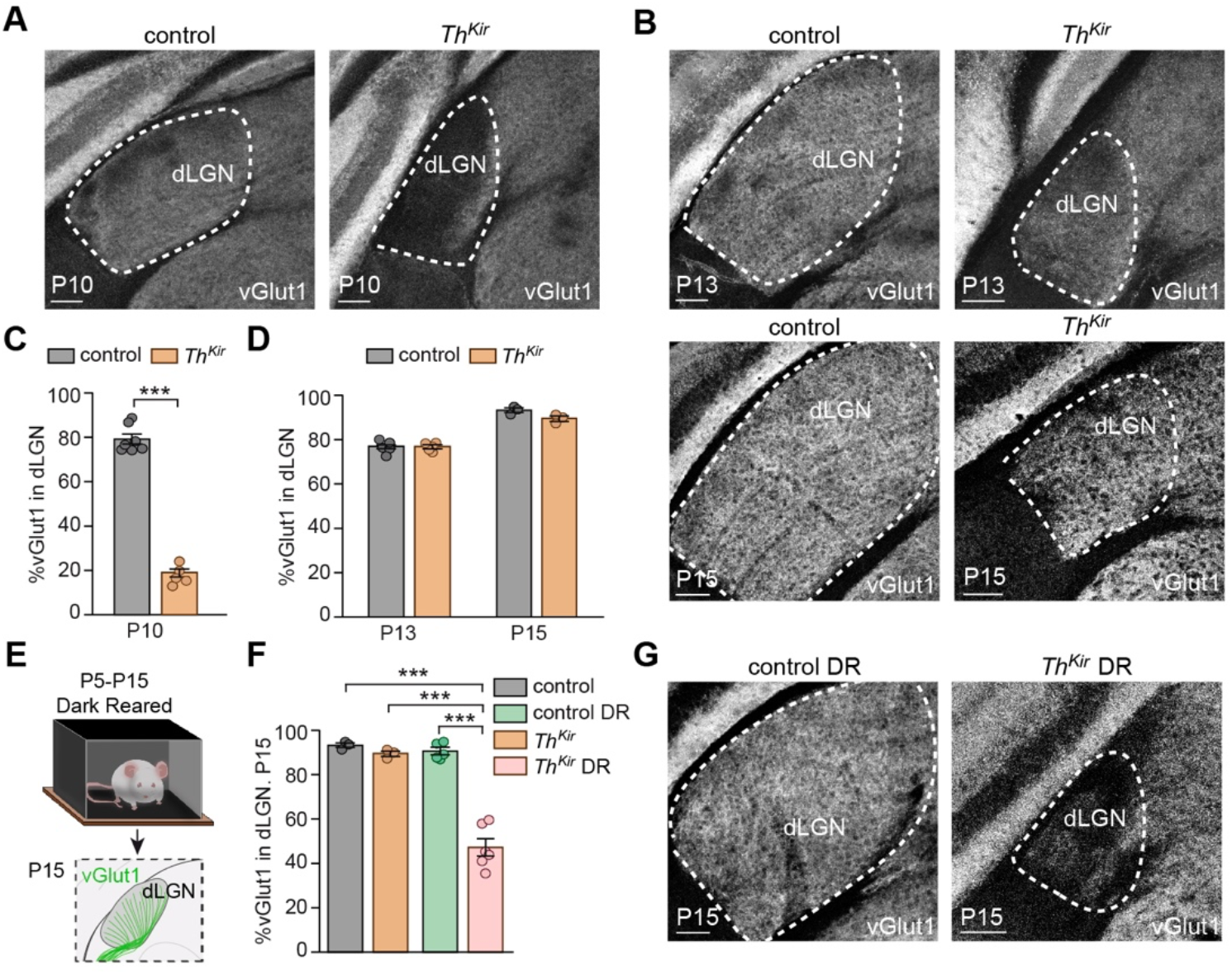
Eye-opening resumes the entry of delayed CTAs into the dLGN of *Th^Kir^* mice. (A) *Left*: Coronal brain sections from control and *Th^Kir^* mice showing vGlut1 immunostaining in the dLGN at P10, before eye opening. (B) *Left*: Coronal sections from control and *Th^Kir^* mice showing vGlut1 immunostaining at P13 and P15, around and after eye-opening, respectively. (C) Quantification of the relative dLGN area occupied by vGlut1 labelling at P10 (control *n* = 7, *Th^Kir^ n* = 6). (D) Quantification of the relative dLGN area occupied by vGlut1 labelling at P13 and P15 (P13: control *n* = 5, *Th^Kir^ n* = 5; P15 control *n* = 3, *Th^Kir^ n* = 3). (E) Scheme of the experimental design used for the dark rearing (DR) experiments. Control and *Th^Kir^* mice were placed in a dark room from P5 to P15, when corticogeniculate innervation was revealed. (F) Quantification of the relative dLGN area occupied by vGlut1 labelling at P15 in mice reared under normal or dark conditions (control *n* = 3, control DR *n* = 5, *Th^Kir^ n* = 3, *Th^Kir^* DR *n* = 6). (G) Coronal sections showing the vGlut1 immunostaining in the dLGN at P15 in mice reared under normal or dark conditions. Graphs represent mean ± SEM, each dot corresponding to a single experimental unit (mouse): *P<0.05, **P<0.01, ***P<0.001. Scale bars, 100 μm.

Visual stimulation through closed eyelids evokes glutamatergic retinal waves and light deprivation in mice from P8 to P14 provokes deficient retinogeniculate segregation (Tiriac et al., 2018), demonstrating that activity driven by light stimulus just prior to eye-opening influences the topographic development of the visual system. Thus, we hypothesized that evoked activity may alter the network properties of dLGN neurons, allowing CTAs to enter the dLGN of *Th^Kir^* mice. To test this possibility, we maintained control and *Th^Kir^* mice in a dark rearing environment from P5 until P15 before analyzing CTA innervation of the dLGN (**Figure 5E**). Dark rearing did not provoke major modifications in CTA innervation in control mice at P15. However, in contrast to the rescue phenotype observed in *Th^Kir^* mice reared under normal conditions, there was significantly less CTA invasion of the dLGN in dark reared *Th^Kir^* mice after eye-opening (**Figure 5F** and **5G**). Together these results demonstrate that early visual experience is necessary for the homeostatic recovery of corticogeniculate innervation in juvenile *Th^Kir^* mice.

## Discussion

Here we describe a novel aspect of the coordination of afferent connectivity to the dLGN by intrinsic thalamic activity. Using a mouse model in which spontaneous activity in the dLGN is disrupted, we have found that CTAs were not able to invade the dLGN in time while RTAs were not affected. The CTA invasion is delayed until eye-opening, when visually evoked activity commences and spontaneous retinal waves subside (Blankenship and Feller, 2010). Visual evoked activity is relevant for this recovery since dark-rearing after eye-opening reduces the delayed CTA innervation to less than 50%.

There are aspects of the development of the corticothalamic connectivity that have been well-characterized, such as its temporal dynamics (including waiting periods), the trajectory of cortical axons throughout the telencephalon and diencephalon, the guidance cues involved, and the specificity between cortical layers and the order of their thalamic targets (Grant et al., 2012; Leyva-Diaz and Lopez-Bendito, 2013). However, other aspects remain elusive, especially those related to the final guidance within the thalamus and the invasion of nuclei.

The neonatal dLGN of mice seems to be a repulsive territory for cortical axons since they wait and accumulate in the ventromedial border of the dLGN until P2-P3 (Jacobs et al., 2007). The extracellular matrix of the dLGN after birth presents high levels of aggrecans, a chondroitin sulfate proteoglycan that normally inhibits axon growth (Brooks et al., 2013). Since L6 neurons express the aggrecan receptor, it is likely that this cue contributes to the neonatal waiting period of cortical axons at the dLGN border. It would be interesting to assess whether the delayed entry of cortical axons into the dLGN observed in *Th^Kir^* mice is related to aggrecan levels. If perturbing the activity pattern upregulates aggrecans, or downregulates proteases that cleave aggrecans (Stanton et al., 2011), the dLGN would remain as a repulsive territory for cortical axons beyond the first postnatal days.

Apart from aggrecans, no other permissive or repulsive guidance molecule has been implicated in the invasion of cortical axons into the dLGN. A comprehensive screening of differentially expressed genes in the developing thalamic nuclei is necessary to begin to understand spatial and temporal sorting of the incoming cortical axons. To this end, the analysis should be focused on genes related to guidance, cytoskeleton remodeling, cell-cell interactions, among others. Using this kind of approach, differentially expressed genes have been discovered comparing the dLGN and the primary auditory nucleus of the thalamus at P5. Some of these genes belong to the signaling pathways of netrins, semaphorins, slits, and ephrins (Horng et al., 2009), suggesting that a different set of guidance factors may contribute to the formation of modality-specific connections. In the thalamocortical system, we have previously shown that activity controls the expression of one of these guidance molecules. The developing thalamocortical projections express Robo1 in an activity-dependent manner and, when Robo1 is downregulated upon Kir2.1 overexpression, the growing axons change their speed of advance (Mire et al., 2012). Thus, it is probable that the activity-dependent modulation of the expression of some, as yet, unknown guidance molecules delays cortical axon entry into the dLGN in the *Th^Kir^* mice. However, with the available evidence, it is not possible to discern whether activity has a permissive or instructive role in this process.

From our experiments conducted in embBE mice, it is clear that activity is not the only or even the main factor controlling cortical axon behavior in the developing thalamus. In accordance with previous publications, removing retinal afferences causes premature entry of cortical axons into the dLGN (Seabrook et al., 2013). This early invasion is partially delayed if the pattern of spontaneous activity is disrupted in the dLGN. The interpretation of these experiments is not straightforward because we are manipulating activity in a model where retinal axons are removed, a fact that already causes abnormal activity. But, the following perspective may help us to understand the results. We stress here that similar conclusions regarding the role of activity in CTA invasion can be drawn comparing both embBE versus embBE/*Th^Kir^* mice or control versus *Th^Kir^* mice. In both cases, we observed that perturbing spontaneous activity delayed the entry of cortical axons, irrespectively of the initial time point of the entry. Thus, we think that these data indicate that there is an activity-dependent contribution.

In any case, our data further supports that a hierarchy exists in the incoming afferences to the dLGN, where retinal axons prevail over cortical axons (Grant et al., 2016). In recent publications, it has been suggested that cortical axons may also modulate initial targeting of the visual thalamus by the RTAs. In these studies, a subpopulation of retinal ganglion cells fails to project to the dLGN in the absence of corticogeniculate innervation (Shanks et al., 2016; Diao et al., 2017). Despite these findings, we showed that the delayed entrance of CTAs did not alter the retinal axons in *Th^Kir^* mice. Indeed, retinal axons reached and invaded the dLGN normally. Thus, retinal axons were not affected by the abnormal behavior of cortical axons or by the abnormal pattern of activity in the thalamus.

The delayed corticogeniculate innervation in *Th^Kir^* mice resumed at eye-opening suggesting that visual experience somehow forces the cortical axons to enter the dLGN. Visual experience plays a pivotal role in the maturation of visual neuronal networks, especially in species such as ferrets and cats (Chapman and Stryker, 1993; Crair et al., 1998; Li et al., 2006). In mice, the introduction of vision has a milder effect on circuit function (Hagihara et al., 2015); for instance, it induces the formation of functional subnetworks and it delays, but not precludes, the maturation of visual acuity and the emergence of critical periods (Kang et al., 2013; Ko et al., 2013). To the best of our knowledge, there is no other scenario in which sensory experience shortly after eye-opening drives complete recovery from a major developmental defect. This is a striking result. Although we did not perform functional assays, we observed an innervation pattern that resembles that of control mice. Further research is needed to elucidate the activity-dependent mechanisms that drive this recovery and to what extent they have an impact on the modulation of ascending visual information.

Finally, we also found that dLGN growth halted when the cortical axons did not enter the nucleus on time. Previous work has shown that enucleation at P0 leads to a decrease in dLGN size and in the gene expression level of ephrin A5 (Kozanian et al 2015). In the *math5* mutant mouse, which represents a model of genetic deafferentation as retinal ganglion cells do not differentiate, the dLGN also shows a decrease in size (El-Danaf et al 2015). However, in this case, dLGN cell density was higher compared with control, suggesting that the reduction in dLGN size does not appear to be a consequence of cell loss. As the size of the dLGN was normal in *Th^Kir^* mice until P3, just before the initial entry of CTAs, it is also possible that CTAs could exert a trophic effect on dLGN cells, so the lack of connections between CTAs and dLGN might be triggering their cell-death. Alternatively, the abnormal pattern of thalamic spontaneous activity might provoke an exacerbated cell-death in the dLGN as has been shown to occur in other brain regions such as the cortex (Blanquie et al., 2017).

In summary, our data indicate that, in addition to retinal input, patterned spontaneous activity in the perinatal thalamus provides a signal that regulates the timing of CTA invasion of the dLGN. These results also provide evidence of an experience-dependent mechanism that helps to recover from early defects in circuits within the visual system.

## Acknowledgments

The authors are grateful to L.M. Rodríguez, B. Andrés and R. Susín for their excellent technical support; to T.Theil and A. Campagnoni for providing the Golli-tau-eGfp mouse line, and to members of G. López-Bendito’s laboratory for stimulating discussions. Funding: this work was supported by grants from the European Research Council (ERC-2014-CoG-647012) and the Spanish Ministry of Science, Innovation and Universities (PGC2018/096631-B-I00, and Severo Ochoa Grant SEV-2017-0723).

## Declaration of Interests

The authors have no competing interests to declare.

## Supplementary Figures

**Figure S1.**
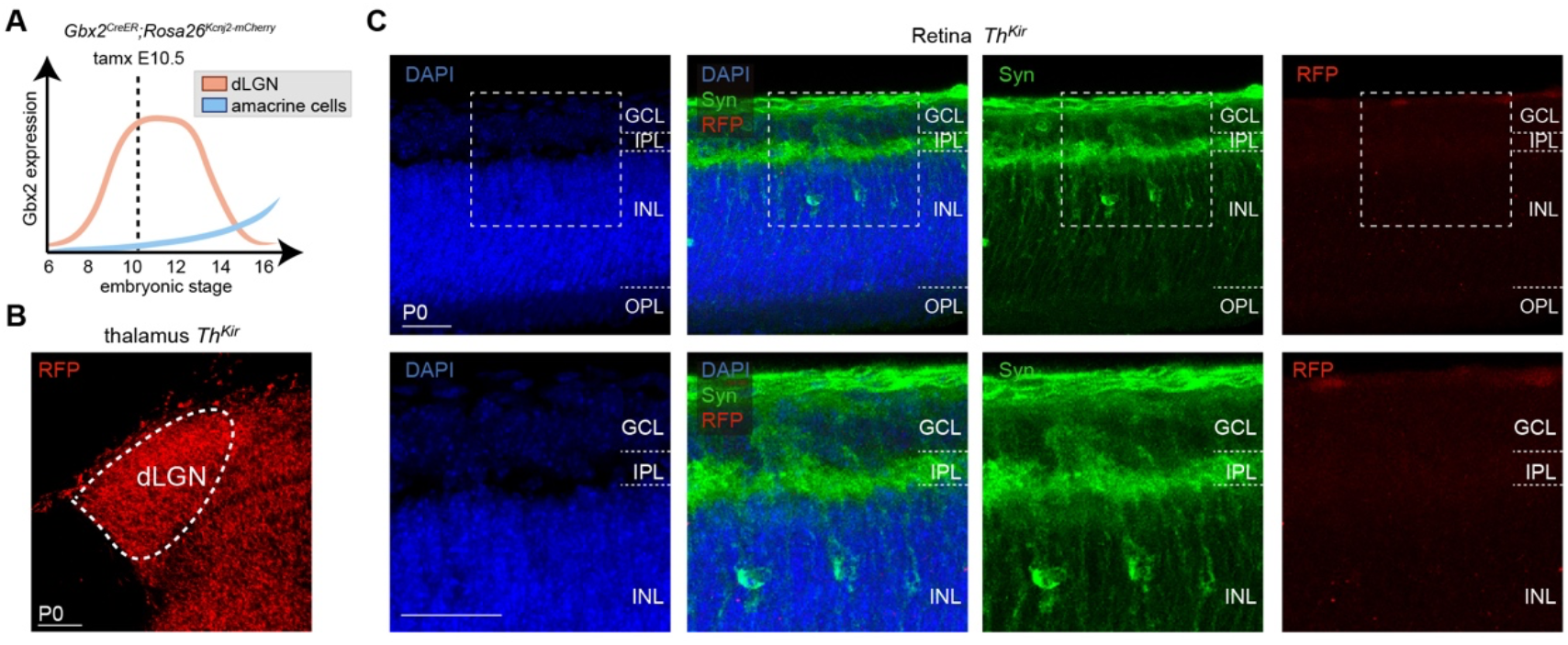
Profile of Kir2.1 expression in the thalamus and retina of *Th^Kir^* mice. (A) Graph depicting the temporal dynamics of the expression of Gbx2 in the dLGN (orange) and retina (blue). Tamoxifen administration at E10.5 selectively targets the dLGN. (B) Red fluorescent protein (RFP) immunostaining in the dLGN and other primary sensory nuclei of the thalamus in coronal brain sections of *Th^Kir^* mice. (C) *Top*, sections of the *Th^Kir^* retina showing that amacrine cells, stained in green against Syntaxin-I (Syn), do not express RFP. GCL corresponds to Ganglion Cell Layer, IPL corresponds to Inner Plexiform Layer, INL corresponds to Inner Nuclear Layer. Retinal layers were outlined using the different cell densities revealed by DAPI staining (control *n* = 6, *Th^Kir^* mice *n* = 6). *Bottom*, magnification of the images above. Scale bars, 100 μm for B and 25 μm for C.

**Figure S2.**
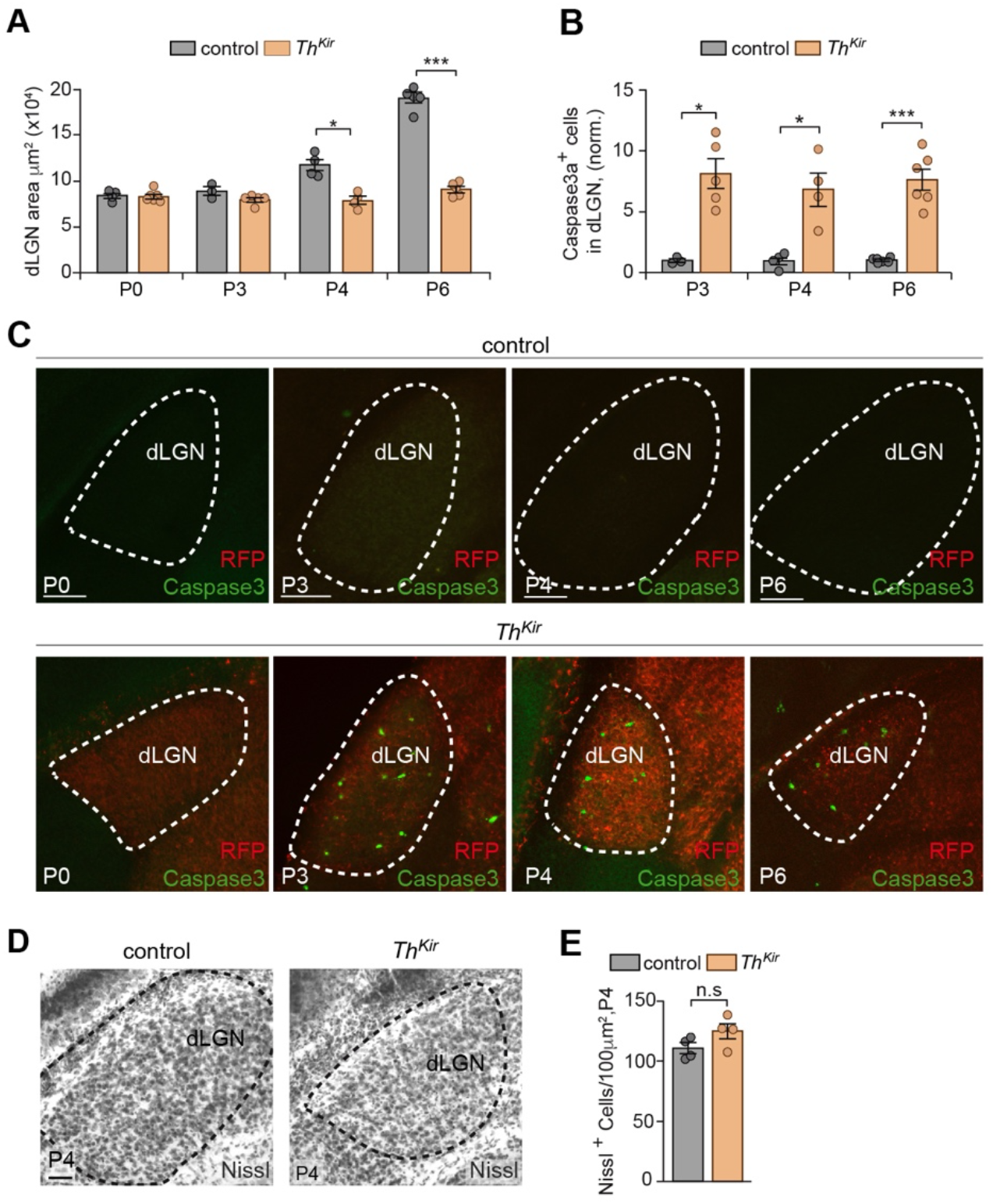
Increased cell-death without changes in the cell density of the dLGN of *Th^Kir^* mice. (A) Quantification of the dLGN area in control and *Th^Kir^* mice (P0 control *n* = 4, *Th^Kir^ n* = 6; P3 control *n* = 3, *Th^Kir^ n* = 5; P4 control *n* = 4, *Th^Kir^ n* = 4; P6 control *n* = 5, *Th^Kir^ n* = 5). (B) Quantification of dLGN cells expressing caspase3a in control and *Th^Kir^* brains at P3, P4 and P6 (P3 control *n* = 3, *Th^Kir^ n* = 5; P4 control *n* = 4, *Th^Kir^ n* = 4; P6 control *n* = 5, *Th^Kir^ n* = 5). (C) Coronal brain sections of control and *Th^Kir^* mice immunostained against caspase3a and RFP (labels Kir2.1-positive cells in *Th^Kir^* mice) at P0, P3, P4 and P6. (D) Nissl stained in coronal brain sections from control and *Th^Kir^* mice at P4. (E) Quantification of the density of Nissl-positive cells in the dLGN (control *n* = 4, *Th^Kir^ n* = 4). Graphs represent mean ± SEM, each dot corresponding to a single experimental unit (mouse): *P<0.05, **P<0.01, ***P<0.001. Scale bars, 100 μm.

**Figure S3.**
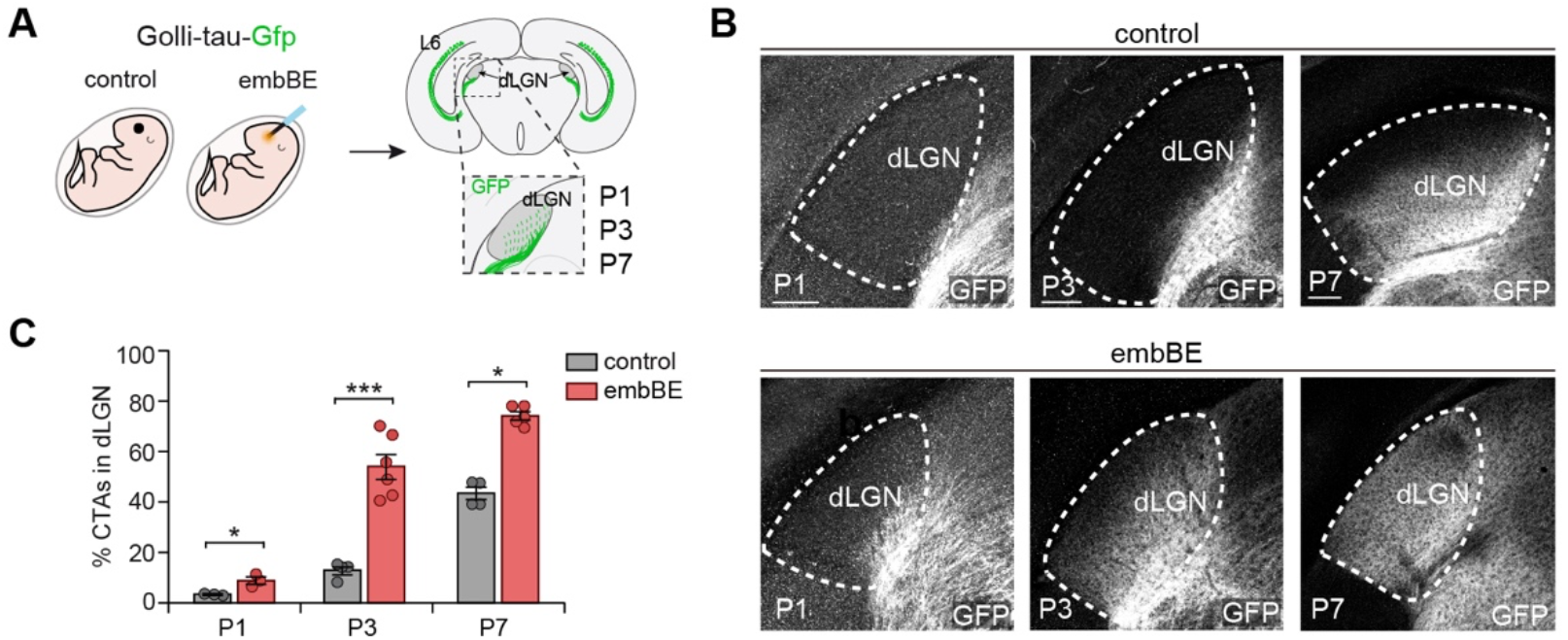
CTAs labeled by Golli-tau-Gfp immunostaining showed a temporal profile of innervation similar to that observed using vGlut1 immunostaining. (A) Scheme of the experimental design used to label cortical axons in Golli-tau-Gfp mice that were enucleated at E14.5 and in their control littermates. (B) GFP immunostaining in coronal sections of brains from control and embBE Golli-tau-Gfp mice at P1, P3 and P7. (C) Quantification of the percentage of the dLGN area occupied by GFP-positive axons (P1 control *n* = 3, embBE *n* = 3; P3 control *n* = 4, embBE *n* = 6; P7 control *n* = 5, embBE *n* = 5). The graph represents mean ± SEM, each dot corresponding to a single experimental unit (mouse): *P<0.05, **P<0.01, ***P<0.001. Scale bars, 100 μm.

**Figure S4.**
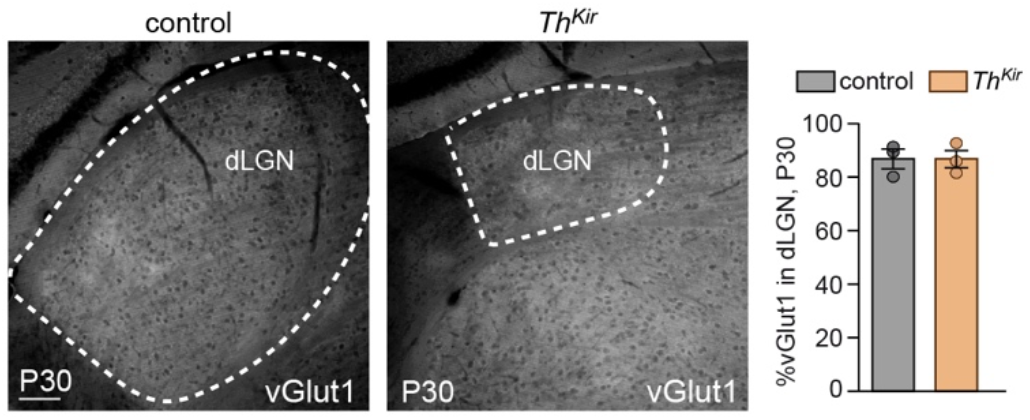
Normal pattern of dLGN invasion by CTAs in adult *Th^Kir^* mice. (A) *Left*, coronal brain sections immunostained against vGlut1 in control and *Th^Kir^* mice at P30. *Right*, quantification of the dLGN area occupied by vGlut1-positive axons at P30 (control *n* = 3, *Th^Kir^ n* = 3), showing no significant differences between both conditions. The graph represents mean ± SEM, each dot corresponding to a single experimental unit (mouse). Scale bar, 100 μm.

## Materials and Methods

### Mouse strains

Wild type mice maintained in an ICR/CD-1 background were used for the *in vivo* experiments, tracing studies and Ca^2+^ imaging. The day on which the vaginal plug was detected was designated as E0.5. The *Rosa26 ^flox-stop (fs)-Kcnj2-mCherry^* mouse line has been described previously (Moreno-Juan *et al*., 2017). This line was crossed with an inducible thalamic-specific *Gbx2^CreER^* line (Brooks *et al*., 2013; Seabrook *et al*., 2013) to generate *Gbx2^CreER/+^; Rosa26 ^fs-Kcnj2^* double mutant embryos, referred to as *Th^Kir^*. Tamoxifen induction of Cre recombination in the double mutant embryos was performed by gavage administration of tamoxifen (5 mg dissolved in corn oil, Sigma) at E10.5 to specifically target the principal sensory thalamic nucleus. The administration of tamoxifen to pregnant mice hinders vaginal delivery. To avoid abortion, we administrated 125 mg/Kg of progesterone (depo-progevera) intraperitoneally (i.p.) at gestational day 14.5 to delay delivery and mice were born via C-section at E19.5. To guarantee safe nursing, we placed newborns with foster mothers. For *Th^Kir^* experiments, CreERT2-negative littermates were used as controls. In the Golli-tau-Gfp mouse line, the golli-promoter of the golli-myelin basic protein gene drives reporter gene expression (enhanced green fluorescent protein (eGFP) fused to bovine tau microtubule protein) selectively in L6 and a subpopulation of L5 cortical neurons, as previously described (Jacobs, 2007). We crossed this line with the *Th^Kir^* animals. Mutants of either sex were analyzed at stages from P0 to P30.

### Histology

For immunohistochemistry, mice were perfused with 4% paraformaldehyde (PFA) in PBS (0.01M), and their brains were removed and post-fixed overnight in the same fixative. For fixation of the retinas, the eyes were dissected out and fixed with 4% PFA for 30 minutes. Immunohistochemistry for all the antibodies was performed on 80-μm sections cut using a vibratome.

Brain sections were first incubated for 1h at room temperature (RT) in a blocking solution containing 1% BSA (Sigma), 2% Donkey Serum (Biowest) and 0.25% Triton X-100 (Sigma) in PBS 0.01M. Subsequently, the sections were incubated overnight at 4 °C with the primary antibodies: guinea-pig anti-vGlut1 (1:5000, Synaptic Systems, #135304), chicken anti-GFP (1:3000; Aves Labs, #Gfp-1020) and rat anti-RFP (1:1000 Chromotek, #5F8), rabbit anti-Caspase3a (1:150; Abcam Ab2302). The sections were then rinsed three times in PBS (0.01M) and incubated for 2h at RT with secondary antibodies: Alexa-546 donkey anti-rabbit (1:500, ThermoFisher, #A10040), Alexa-488 goat anti-chicken (1:500, ThermoFisher, #A11039), Alexa-594 donkey anti-rat (1:500, ThermoFisher, #A21209), Alexa-488 donkey anti-rabbit (1:500, ThermoFisher, #A21206), Alexa-647 donkey anti-Guinea Pig (1:500, ThermoFisher). Finally, the sections were counterstained with the fluorescent nuclear dye DAPI (Sigma-Aldrich).

Retinal sections were first incubated for 30 min at 37 °C in a 2N HCl solution for antigen retrieval. After two 5-minute washes in 0.1M sodium borate (pH 8.5), the slices were incubated at RT for 1 hour in a blocking solution containing 3% Donkey Serum (Biowest) and 0.3% Triton X-100 (Sigma) in PBS 0.01M. Subsequently, the sections were incubated overnight at 4 °C with the primary antibodies: mouse anti-Syntaxin-1 (1:500, Sigma, # S0664), chicken anti-GFP (1:3000; Aves Labs, #Gfp-1020) and rat anti-RFP (1:1000 Chromotek, #5F8). The sections were then rinsed three times in PBS (0.01M) and incubated for 2 hours at RT with secondary antibodies: Alexa-647 donkey anti-mouse (1:500, Mol. Probes A31571, #A10040), Alexa-488 goat anti-chicken (1:500, ThermoFisher, #A11039), Alexa-594 donkey anti-rat (1:500, ThermoFisher, #A21209). Finally, the sections were counterstained with the fluorescent nuclear dye DAPI (Sigma-Aldrich). For Nissl staining, the 80-μm sections were mounted on slides and dried overnight. The slides were incubated in a cresyl violet solution for 30 seconds to 2 minutes, dehydrated in an ethanol series (70% ethanol for 3 minutes, 70% ethanol for 3 minutes, 90% ethanol for 3 minutes, 90% ethanol for 3 minutes, 100% ethanol for 3 minutes), passed through Xylol (2 x 5 minutes) and immediately mounted with Eukitt^®^ (ThermoFisher).

### Viral tracing of visual corticothalamic axons

For cortical axonal tracing at postnatal day 9, a lentivirus containing Gfp (pLM-CMV-tauEGFP) was injected into the V1 cortex of control and *Th^Kir^* mice at P3. Pups were anesthetized under hypothermia and 5 pulses of 69 nL were injected into the V1 with a nanoliter injector (WPI). Four days later, the animals were perfused with 4% PFA in PBS (0.01M) and their brain was dissected out and post-fixed overnight in the same fixative solution. Vibratome sections (thickness: 80 μm) were counterstained with DAPI (Sigma-Aldrich).

### Tracing of retinal axons

For axonal tracing at postnatal stages, P0 pups were anesthetized under hypothermia and P6 pups were anesthetized with isoflurane. An incision was made over the eyelid of one eye and it was injected with CTB conjugated to Alexa-647 (Mol. Probes C34778) using a nanoliter injector (WPI). The animals were perfused with 4% PFA in PBS (0.01M) 2, 3, or 7 days later, and their brain was dissected out and post-fixed overnight in the same fixative. Vibratome sections (thickness: 80 μm) were then counterstained with the fluorescent nuclear dye DAPI (Sigma-Aldrich).

### *In utero* enucleation

For *in utero* enucleation, wild type or *Th^Kir^* pregnant females (gestational day 14.5) were deeply anesthetized with isoflurane to perform laparotomies. The embryos were exposed and both eyes were cauterized in half of the litter. The surgical incision was closed and the embryos were allowed to develop until postnatal stages.

### Calcium imaging in slices

The brains of postnatal mice (P0 and P1) obtained after they were sacrificed by decapitation were used for calcium imaging experiments. The brains were immediately dissected out and 300-μm thick coronal slices were cut using a vibratome (VT1200 Leica Microsystems Germany) using a slicing solution bubbled with 95% O_2_ and 5% CO_2_ and containing (mM): 2.5 KCl, 7 MgSO_4_, 0.5 CaCl_2_, 1 NaH_2_PO_4_, 26 NaHCO_3_, 11 glucose and 228 sucrose. The slices were kept at RT in oxygenated artificial CSF (aCSF) containing (mM): 119 NaCl, 5 KCl, 1.3 MgSO_4_, 2.4 CaCl_2_, 1 NaH_2_PO_4_, 11 glucose. To load the calcium indicator, the slices were incubated at 37 °C for 30 minutes in oxygenated aCSF containing 5 μl of Cal520 AM (AAT Bioquest), 1 mM DMSO and 20% pluronic acid. The slices were then placed under a Leica DMi8 microscope in a recording chamber perfused with oxygenated aCSF at 34 °C. We acquired images every 300 ms for 15 minutes using a Hamamatsu ORCA-Flash 4.OLT camera. A minimum of three timelapse movies were obtained per slice. We used one slice per animal and each data point in the graph corresponds to one independent observation. Calcium imaging data were analyzed as extensively described elsewhere (Moreno-Juan *et al*., 2017; Antón-Bolaños *et al*., 2019). Briefly, the dLGN was delineated in each slice and this area was parcellated into ROIs using a grid of 6×6 pixel squares. Each ROI had the approximate size of a cell. Calcium events were detected in each ROI and the fraction of the total ROIs active was quantified throughout the movie.

### Dark rearing experiments

For the DR experiments, *Th^Kir^* pups and their control littermates were housed into a completely dark environment with their lactating dam from P5 to P15. After this, the pups were perfused with 4% PFA in PBS (0.01M). Their brain was dissected out and postfixed overnight in the same fixative. Vibratome sections (thickness: 80 μm) were then subjected to immunohistochemistry.

### Data analysis and statistics

The analysis of the calcium imaging experiments was described in detail elsewhere (Moreno-Juan *et al*., 2017; Antón-Bolaños *et al*., 2019). In order to delineate the dLGN we excluded the optic tract and we followed the clear separation of the dLGN with the ventral lateral geniculate nucleus and the ventral posterior medial nucleus. ImageJ software was used to quantify CTA and RTA invasion of the dLGN. Coronal slices (thickness: 80 μm) were obtained from the animals subjected to the different conditions and immunostained against vGlut1 or GFP. At least three dLGN sections were analyzed from each animal, correcting for the fluorescence background. Threshold fluorescence images were analyzed using ImageJ, analyzing the particles of the pixels that cover the dLGN and measuring the proportion of pixels occupied by CTAs. For the analysis, we used at least three slices from each animal calculating the percentage of innervation. The average value of the three sections was considered an independent observation.

Caspase3a positive cells were quantified in coronal sections (thickness: 80 μm) and normalized to the area of each dLGN. The whole dLGN was measured from P3, P4 and P6 control and *Th^Kir^* mice, and the results normalized to the mean of the controls, representing the fold-change in Caspase3a positive cells in the *Th^Kir^* mice.

Statistical analysis was carried out with GraphPad Prism6™ and Matlab™, and the data are presented as mean and SEM. Statistical comparison between the groups was performed using an unpaired two-tailed Student’s *t* test and when the data failed a Kolmogorov-Smirnov or a Shapiro Wilk test of normality, a two-tailed, non-parametric Mann-Whitney U-test was used. A one-way ANOVA test was performed on the embryonic enucleation in *Th^Kir^* mice experiments. A two-way ANOVA test was performed on the embryonic enucleation in *Th^Kir^* mice and for the DR experiments. No statistical methods were used to predetermine the sample size, although our sample sizes are considered adequate for the experiments and consistent with the literature (Brooks *et al*., 2013; Seabrook *et al*., 2013; Grant *et al*., 2016). The allocation of mice to the experimental groups was not randomized.

### Quantifications

#### Figure 1

Figure 1D Event frequency (events/minute): control(1-50%) 0.29 ± 0.05, control(50-100%) 0.043 ± 0.008, *n* = 10; *Th^Kir^*(1-50%) 0.18 ± 0.09, *Th^Kir^*(50-100%) 0 ± 0, *n* = 4 (P = 0.31, two-tailed student’s *t*-test; **P 0 0.0044, two-tailed student’s *t*-test). Figure 1E Total event frequency (event/minute/ROI): 0.14 ± 0.014, *n* = 10; *Th^Kir^* 0.036 ± 0.015, *n* = 4 (*P = 0.017, two-tailed student’s *t*-test).

#### Figure 2

Figure 2C vGlut1 at P4: control 10 ± 1%, *n* = 5; *Th^Kir^* 1.67 ± 0.34%, *n* = 5 (***P<0.001, two-tailed student’s *t*-test). At P6: control 35.96 ± 4.47%, *n* = 5; *Th^Kir^* 1.24 ± 0.51%, *n* = 5 (***P<0.001, two-tailed student’s *t*-test). At P9: control 62.4 ± 4.49%, *n* = 5; *Th^Kir^* 12.15 ± 0.74%, *n* = 3 (***P<0.001, two-tailed student’s *t*-test). Figure 2G Percentage of RTAs in dLGN at P2. Contralateral RTAs in dLGN, control: 69.94 ± 1.56%, *n* = 3; *Th^Kir^*: 66.5 ± 1.74%, *n* = 4. P=0.22, two-tailed student’s *t*-test. Ipsilateral RTAs in dLGN control: 20.45 ± 2.27%, *n* = 3; *Th^Kir^*: 24.34 ± 2.3%, *n* = 4 (P = 0.29, two-tailed student’s *t*-test).

#### Figure 3

Figure 3C % of vGlut1 in the dLGN at P2: control 7.1 ± 0.92%, *n* = 3; embBE 44.32 ± 2.35%, *n* = 3 (***P<0.001, two-tailed student’s t-test). At P3: control 10.68 ± 1.92%, *n* = 3; embBE 53.06 ± 4.41%, *n* = 3 (***P<0.001, two-tailed student’s t-test). At P7: control 39.01 ± 3.03%, *n* = 3; embBE 71.45 ± 1.11%, *n* = 3 (***P<0.001, two-tailed student’s t-test). Figure 3F Event frequency (event/minute): control(1-50%) 0.20 ± 0.04, control(50-100%) 0.039 ± 0.02, *n* = 9; embBE(1-50%) 0.19 ± 0.03, embBE(50-100%) 0.11 ± 0.03, *n* = 5 (P = 0.77, two-tailed student’s *t*-test; *P = 0.017, two-tailed student’s *t*-test). Figure 3H Event frequency (event/minute): control(1-50%) 0.40 ± 0.11, control(50-100%) 0.48 ± 0.48, *n* = 8; embBE(1-50%) 0.39 ± 0.18, embBE(50-100%) 0.18 ± 0.06, *n* = 6 (P = 0.98, two-tailed student’s *t*-test; **P = 0.0096, two-tailed student’s *t*-test).

#### Figure 4

Figure 4C Event frequency at P0 (event/minute): embBE(1-50%) 0.19 ± 0.03, embBE(50-100%) 0.11 ± 0.03, *n* = 7; embBE *Th^Kir^* (1-50%) 0.09 ± 0.03, embBE *Th^Kir^* (50-100%) 0.0 ± 0.0, *n* = 4 (P = 0.07, two-tailed student’s *t*-test; *P = 0.011, two-tailed student’s *t*-test). Figure 4F % of vGlut1 in the dLGN at P7: control *n* = 5, embBE *n* = 5, embBE *Th^Kir^ n* = 4. A one-way ANOVA test with Tukey’s post hoc analysis indicates that the interaction between all the conditions are significant, comparisons: control vs embBE *** P<0.001; control vs embBE *Th^Kir^* *P = 0.01; embBE vs embBE *Th^Kir^* **P = 0.0026.

#### Figure 5

Figure 5C % vGlut1 in the dLGN of P10: control 79.28 ± 2.31%, *n* = 7; *Th^Kir^* 18.85. ± 1.85%, *n* = 6 (***P<0.001, two-tailed student’s *t*-test). Figure 5D % vGlut1 in the dLGN of P13: control 76.62 ± 1.23%, *n* = 5; *Th^Kir^* 76.69. ± 0.8%, *n* = 5 (P = 0.96, two-tailed student’s *t*-test) and % vGlut1 in the dLGN at P15: control 93.17 ± 1%, *n* = 3; *Th^Kir^* 89.31 ± 1.16%, *n* = 3 (P= 0.07 two-tailed student’s *t*-test). Figure 5F % vGlut1 in the dLGN at P15: control *n* = 3, control DR *n* = 5, *Th^Kir^ n* = 3, *Th^Kir^* DR *n* = 6. A two-way ANOVA test with Tukey’s post hoc analysis, comparisons: control vs *Th^Kir^* P = 0.97; control vs control DR P = 0.99; control vs *Th^Kir^* DR ***P<0.005; *Th^Kir^* vs control DR P = 0.99; *Th^Kir^* vs *Th^Kir^* DR ***P<0.001; control DR vs *Th^Kir^* DR ***P<0.001.

#### Figure S2

Figure S2A dLGN size at P0: control 84326 ± 2872, *n* = 4; *Th^Kir^* 83720 ± 2498, *n* = 6 (P = 0.88, two-tailed student’s *t*-test). At P3: control 89800 ± 4558, *n* = 3; *Th^Kir^* 80403 ± 2300, *n* = 5 P = 0.14, Mann-Whitney *U*-test. P4: control 118410 ± 6262, *n* = 4; *Th^Kir^*: 79748 ± 4351, *n* = 4 (*P = 0.02, two-tailed student’s *t*-test). At P6: control 193136 ± 5603, *n* = 5; *Th^Kir^* 92022 ± 3380, *n* = 5 (***P<0.001, two-tailed student’s *t*-test). Figure S2B Caspase3a^+^ cells normalized to the control data at P3: control 100 ± 14.97%, *n* = 3; *Th^Kir^* 816.3 ± 122.3%, *n* = 5 (*P = 0.03, Mann-Whitney *U*-test). At P4 control 100 ± 30.98%, *n* = 4; *Th^Kir^* 684.4 ± 139.2%, *n* = 4 (*P = 0.028, Mann-Whitney *U*-test). At P6 control 100 ± 7.71%, *n* = 6; *Th^Kir^* 762.6 ± 87.69%, *n* = 6 (***P<0.001, two-tailed student’s *t*-test). Figure S2E Nissl^+^ cells/100 μm^2^: control 110.5 ± 4.29, *n* = 4; *Th^Kir^* 124.8 ± 6.34%, *n* = 4 (P = 0.11. Mann-Whitney *U*-test).

#### Figure S3

Figure S3C % CTAs in the dLGN at P1: control 3.26 ± 0.2%, *n* = 3; embBE 8.85 ± 1.44%, *n* = 3 *P = 0.018, two-tailed student’s t-test). At P3: control 12.86 ± 1.59%, *n* = 4; embBE 53.92 ± 5.05%, *n* = 6 (***P<0.001, two-tailed student’s t-test). At P7: control 43.89 ± 2.46%, *n* = 4; embBE 74.82 ± 1.68%, *n* = 5 (*P = 0.01, Mann-Whitney U-test).

#### Figure S4

Figure S4 % vGlut1 at P30: control 87.26 ± 3.36%, *n* = 3; *Th^Kir^* 87.16. ± 3.28%, *n* = 3 (P = 0.98, two-tailed student’s *t*-test).

## Notes

### Competing Interest Statement

The authors have declared no competing interest.

